# Cost of altered translation accuracy shapes adaptation to antibiotics in *E. coli*

**DOI:** 10.1101/2025.06.28.662112

**Authors:** Laasya Samhita, Sharvari Tamhankar, Joshua Miranda, Aabeer Basu, Deepa Agashe

## Abstract

Protein synthesis, while central to cellular function, is error-prone. The resulting mistranslation is generally costly, but we do not know how these costs compare or interact with the costs imposed by external selection pressures such as antibiotics. We also do not know whether and how these costs are compensated during evolution. It is important to answer these questions, since mistranslation is ubiquitous and antibiotic exposure, widespread. We quantified the growth cost of genetically increasing and decreasing mistranslation rates, and exposure to low antibiotic concentrations, in *Escherichia coli*. Mistranslation costs were generally lower than the cost imposed by antibiotics, and exacerbated in a strain-specific manner under antibiotic exposure. All strains quickly compensated the antibiotic cost during experimental evolution, via antibiotic- and genotype-specific mutations. In contrast, mistranslation costs were significantly reduced only in some cases, without clear causal mutations. Control populations that evolved without antibiotics consistently compensated the cost of accuracy, and evolved increased antibiotic resistance as a by-product. Our work demonstrates that even when the cost of mistranslation is weak, altered translation accuracy can shape adaptive outcomes and underlying genetic mechanisms, with strong collateral fitness effects for apparently unrelated phenotypes such as antibiotic resistance.

## Introduction

Mistranslation contributes to non-genetic variation in all life forms. Translation errors and the resultant heterogenous protein sequences were long considered sources of cellular noise, but it is now clear that they have significant, sometimes even predictable, phenotypic effects. Mistranslation is typically costly (Mohler and Ibba 2017), reducing mean division times at the single cell level, and lowering growth rates at the population level (Samhita et al., 2021). Such costs are thought to be mediated primarily by misfolding or aggregation of mistranslated proteins (Drummond and Wilke 2008, Paredes et al., 2012) as well as reduced availability of functional proteins (Ruan et al., 2008, Bullwinkle and Ibba 2016). These broad costs also mean that higher levels of global mistranslation can exacerbate the effect of deleterious mutations (Bratulic et al., 2017, Schmutzer and Wagner 2023) or reduce the fitness advantage of beneficial mutations (Schmutzer and Wagner 2023). How do cells prevent such mistranslation or reduce its fitness effects? Across evolutionary timescales, evidence from comparative genomics indicates that cost mitigation can occur via multiple mechanisms. For example, in diverse taxa, protein coding sequences preferentially use codons that minimize mistranslation, and this pattern gets stronger with increasing gene expression (Drummond and Wilke 2008). Simulations suggest that normally untranslated sequences in the 3’UTR that are occasionally read through (leading to mistranslation) are selectively purged (Kosinski and Masel 2020). Despite these cost-minimizing strategies, mistranslation can still present a significant cellular burden. Under these circumstances, over-expressing chaperones or increasing protease levels can immediately buffer the phenotypic effects of mistranslation (Ruan et al., 2008, Samhita et al., 2020), presumably by decreasing the load of non-functional and aggregated proteins. Thus, there is substantial evidence for the deleterious consequences of mistranslation, and preventive and compensatory mechanisms to combat it.

On the other hand, a growing body of work also shows a range of benefits of mistranslation (Javid et al., 2014, Schwartz et al., 2016, Samhita et al., 2020), especially under external selection pressures that also impose fitness costs. For example, mistranslating *Escherichia coli* cells under heat stress show higher survival than their WT counterparts (Evans et al., 2019). Under oxidative stress, both eukaryotic and bacterial cells increase the incidence of a particular form of mistranslation — specific tRNA misacylation — with a concomitant survival benefit (Netzer et al., 2009, Schwartz et al., 2016). The effect of mistranslation on the response to antibiotic exposure has been a particular favourite, explored across several studies (Javid et al., 2014, Bratulic et al., 2015, Samhita et al., 2020). Mistranslation can lead to faster adaptation, a higher frequency of antibiotic resistance as compared with the wild type (Javid et al., 2014, Weil et al., 2017, Samhita et al., 2020), or different mutational outcomes (Zheng et al., 2021). Mistranslation can also confer antibiotic resistance via novel mechanisms that are as yet unclear (Schwartz et al., 2016). In contrast, some studies comparing mistranslating vs. wild type *E. coli* either found no difference in the rate or effect of mutations in antibiotic resistance genes (Bratulic et al., 2017), or found identical mutations under selection from inhibitory antibiotic concentrations (Samhita et al., 2020). Thus, although mistranslation can have both positive and negative impacts for cells, our ability to predict net outcomes – especially across evolutionary timescales – remains poor.

One reason for the poor predictive power is that we do not understand exactly how the costs imposed by mistranslation compare or interact with the costs imposed by external stressors such as antibiotics. This is important, because these interactions may determine possible adaptive mechanisms, ultimately influencing evolutionary outcomes. For example, in the *E. coli* experiments mentioned above, under strong selection (high antibiotic dose), mistranslating populations fixed amino acid changes that stabilized protein structure; while under weak selection (low antibiotic dose) a reduction in protein expression appeared to drive adaptation (Bratulic et al., 2015). In later work, this effect was attributed to epistasis between the genetic background (which determined the level of mistranslation) and the phenotypic effect of new mutations (Zheng et al., 2021). Higher levels of mistranslation generally exacerbated the negative impact of deleterious mutations, altering the identity of fixed mutations during evolution (Bratulic et al., 2017). However, the observed fitness variation across strains could not be separately attributed to costs imposed by mistranslation versus antibiotics, or compensation thereof. This is especially important to delineate with antibiotic selection because cost compensation for a given antibiotic can vary with genetic background, study design (*in vitro/in vivo*), drug concentration, and the temporal context of antibiotic treatment (e.g., administering different antibiotics sequentially or in combination) (reviewed in Andersson and Hughes 2010). Thus, it is unclear if the costs of antibiotic change with mistranslation levels, or whether the two costs interact to influence adaptive dynamics under antibiotic selection. One study that inadvertently combined both costs involved the antibiotic tavabarole. This synthetic antibiotic inactivates the essential enzyme leucine tRNA synthetase (LeuRS) that amino-acylates tRNA^Leu^. Like most synthetases, LeuRS contains an editing domain that removes incorrectly incorporated amino acids, increasing translation accuracy. The binding site of tavabarole falls within the editing domain, such that resistance-conferring mutations often increase mistranslation (Melnikov et al., 2020). As a result, such mutants develop antibiotic resistance while simultaneously incurring a cost of mistranslation. Overall, it is clear that costs of mistranslation and antibiotic exposure could interact in multiple ways. Separating these costs, and their interactions, is necessary to understand how the costs vs benefits of mistranslation may alter the dynamics of adaptation to antibiotics.

To understand the impact of both mistranslation and antibiotic exposure on adaptive trajectories, we simultaneously quantified the costs of altered translation accuracy and cost of exposure to antibiotics in *E. coli*. From previous work, we know that both increasing and decreasing translation accuracy can lead to a fitness cost (Samhita et al., 2021). We therefore evolved strains with both higher and lower translation accuracy than the wild type strain (WT), in sub-inhibitory concentrations of three antibiotics with differing modes of action: ciprofloxacin (causes DNA damage), ampicillin (blocks cell wall biosynthesis) and kanamycin (protein synthesis inhibitor, also causes mistranslation). Using growth rate measurements of ancestral and evolved lines, we defined the ancestral costs of antibiotic and altered translation accuracy and investigated how these costs changed during the course of evolution at both the phenotypic and genotypic levels **(Figure 1)**. We find that while antibiotic costs dominate and are rapidly compensated, the costs of altered accuracy are compensated in specific genotype by environment combinations. We previously reported that both WT and mistranslating cells fix the same beneficial mutations during short-term adaptation to inhibitory concentrations (~5X MIC, minimum inhibitory concentration) of ciprofloxacin (Samhita 2020). Here, under milder selection pressures (sub-MIC) imposed over a longer time period, we find mutational diversity across strains in all antibiotics, suggesting that evolutionary predictability is confounded both by altered accuracy and antibiotic selection at sub-MIC.

**Figure 1.**
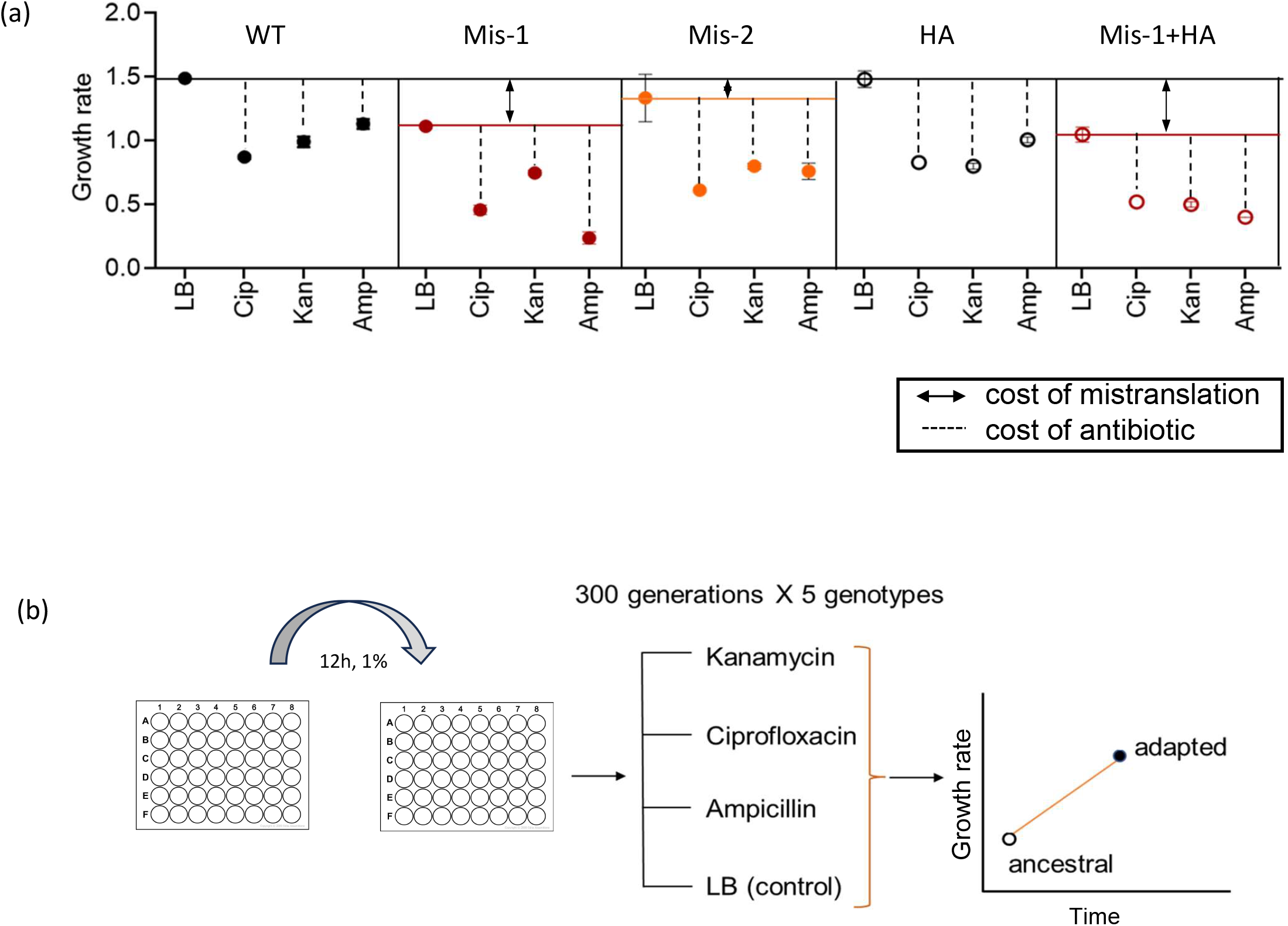
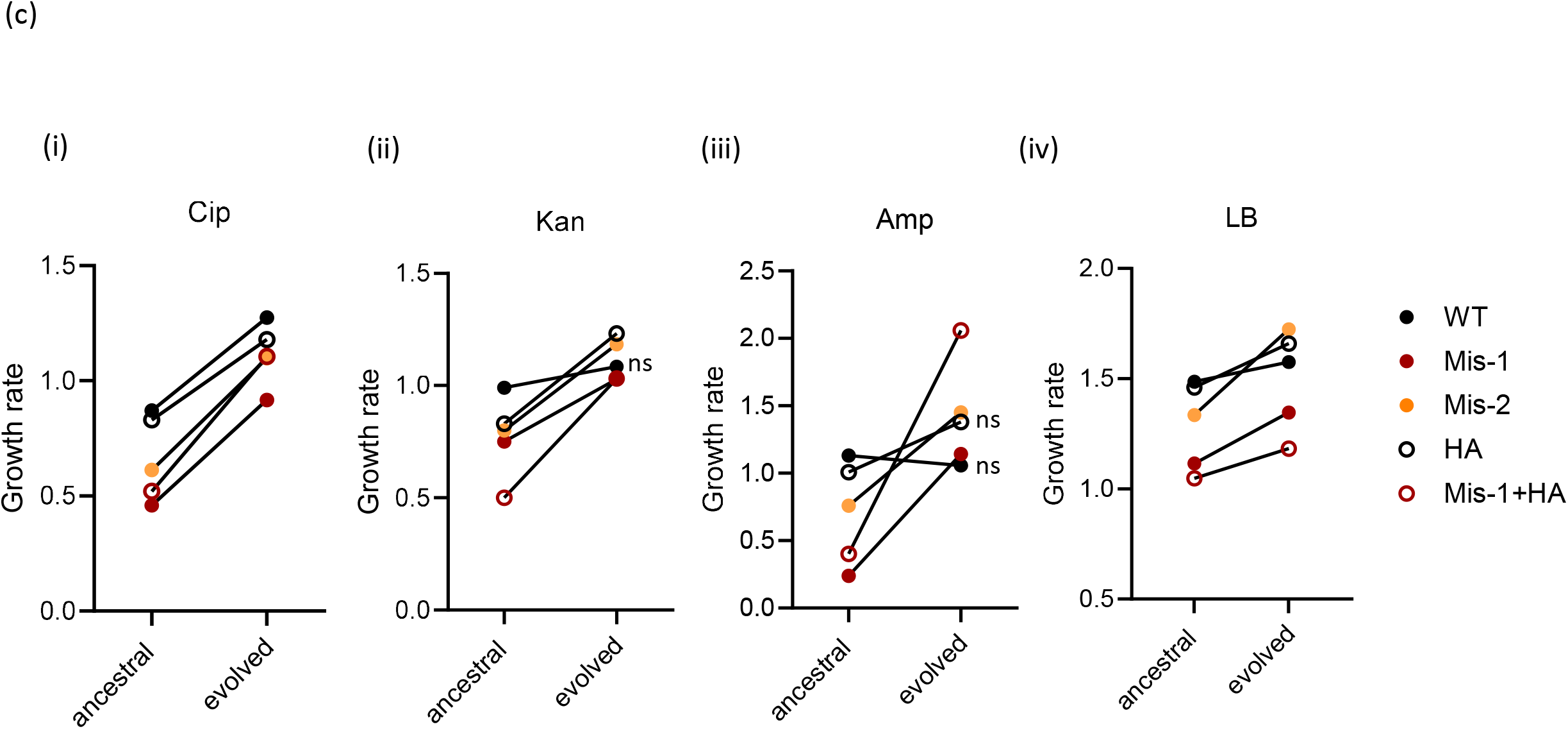
**(a) Ancestral costs of accuracy and exposure to antibiotic.** Mean growth rates (+/−SD) of all five ancestral strains in the four growth regimes used in this study. Dotted lines indicate antibiotic cost while double-headed arrows indicate accuracy cost (n=5 for all except WT and Mis-1 in Kan, Cip and LB, where n=6 and HA in LB, where n=4). Note that the figure shows growth rate differences rather than ratios which we use in computing cost values (See Methods). **(b) Schematic for experimental evolution**. We initiated cultures for our experimental evolution regime using single colonies from an agar plate (n=5 for all except except WT and Mis-1 in Kan, Cip and LB, where n=6 and HA in LB, where n=4). We grew cultures in 48 well plates under shaking conditions at 37°C in four regimes: LB with one of three antibiotics, and LB alone (control). We assessed adaptation using change in maximum growth rate. **(c) Most strains rapidly adapt to low antibiotic concentrations across regimes**. Comparison of mean growth rates for ancestral vs evolved lines, tested in the same regimes in which they evolved (n=5 for all except except WT and Mis-1 in Kan, Cip and LB, where n=6 and HA in LB, where n=4). All lines showed a significant increase in growth rate except when indicated (‘ns’, not significant; Table S1).

## MATERIALS AND METHODS

### Bacterial strains

We used KL16 *E. coli* cells (Low 1968) as our wild type strain (WT). To generate strains with altered accuracy, we used four genetically altered derivatives of the WT: two mistranslating (Mis-1 and Mis-2), one hyper-accurate (HA) and one with a combination of a mistranslating and hyper-accurate strains (Mis-1+HA). The KL*ΔZWV* strain (‘Mis-1’), lacks three of four initiator tRNA genes encoded by *E. coli* (Samhita et al., 2013). Initiator tRNA acts only at the first step of protein synthesis and has no substitute (Gualerzi and Pon 2015). This strain therefore has a lower rate of protein synthesis, a ~20% slower growth rate than the WT (Samhita et al., 2020), and mistranslates through at least one mechanism: initiation of translation at non-AUG codons (Samhita et al., 2013). The KL*rpsDE68D* strain (‘Mis-2’) mistranslates by reducing the fidelity of codon-anticodon interactions on the ribosome, providing us with an independent mechanism of mistranslation (Agarwal et al., 2015). To increase translation accuracy in the WT, we introduced a mutation (K42R) in the ribosomal protein S12, generating strain KL*rpsLK42R* (‘HA’ for ‘hyper-accurate’). The mutation reduces the frequency of errors during mRNA decoding (Chumpolkulwong et al., 2004). Finally, we introduced the same mutation into strain Mis-1, generating the hybrid strain ‘Mis-1+HA’. Note that the introduction of hyper-accurate ribosomes does not reverse the non-AUG initiation seen in Mis-1, however it lowers overall background mistranslation levels by reducing decoding errors (Samhita et al., 2020).

### Growth conditions and media

We grew bacterial cultures in Luria Bertani medium (LB) in 48-well plates under shaking conditions or on LB-agar plates containing 1.8% (w/v) agar (Difco) in static conditions, in both cases at 37°C. In the experimental evolution regime, we added antibiotics (Sigma-Aldrich) at the following sub-inhibitory concentrations: ampicillin (amp) at 2.25 µg/mL (MIC for WT KL16 is ~4 µg/mL), kanamycin sulphate (Kan) at 4 µg/mL (MIC ~10 µg/mL), ciprofloxacin (Cip) at 10 ng/mL (MIC ~20 ng/mL) and kasugamycin (Ksg) at 30 µg/mL (MIC ~ 10 µg/mL). Kasugamycin inhibits initiation of protein synthesis (Schluenzen et al., 2006), and served as a tRNA-independent method of reducing translation initiation events, i.e., a control for our mistranslating line which lacks three of four initiator tRNA genes. In all cases, the exact sub-MIC antibiotic concentration used in the experiments was set by the concentration beyond which the Mis-1 genotype (the slowest strain) either did not grow or showed severe growth retardation within 12 h.

### Experimental evolution under antibiotic selection

Bacterial populations were grown in plain LB or LB supplemented with one of three antibiotics as described **(Fig. 1b)**, generating 4 treatments per strain with 5-6 biological replicates per strain per treatment (five well-separated independent colonies from a freshly streaked plate of each strain). Populations were grown in 48-well plates for ~300 generations (15 days) under shaking conditions at 37°C. Additionally, we also evolved WT supplemented with kasugamycin to serve as a control. All lines were sub-cultured every 12 h, with 1% by volume transferred to fresh medium (5 µL in 500 µL). Every 3 days and finally on the 16^th^ day, stocks of the evolved (mixed) cultures were stored in 40% glycerol at −80°C for further analysis. Due to practical constraints, WT and Mis-1 were evolved first across all regimes, followed by HA and Mis-1+HA and finally Mis-2 and WT(Ksg), in three separate sets with an overall time period of 3 months.

### Measuring growth rate and MIC as a fitness proxy

We measured growth rates of evolved populations and their respective ancestors in the regime they were evolved in, as well as in LB. Conversely, lines evolved in LB were also tested for growth in antibiotic-containing media. We first inoculated a loopful of each freezer stock in LB for ~8 h, and then transferred 5 µL to 495 µL of LB with antibiotics (as needed) at the same concentration they were subjected to during the experimental evolution, and allowed cultures to grow at 37 °C with shaking at 200 rpm for 12-14 h. We inoculated 5 µL of these pre-cultures into 495 µL growth media in 48-well plates (Corning) and incubated in an automated growth reader (Biotek), with shaking, at 37 °C, recording absorbance at O.D_600nm_ every 15 minutes for 18-24 h. We measured the growth rate of three technical replicates per biological replicate in each growth medium. In each 48-well plate, we included the relevant ancestors in the same medium to estimate and account for day-to-day variation in growth rates. Note that ancestral values did not in fact vary significantly and the same 5 biological replicates are represented in each figure. Maximum growth rate in early log phase (below OD_600nm_ ~ 0.3) was estimated using the Curve Fitter software (Delaney et al. 2013). We compared ancestral with evolved growth rates and used the value as a proxy for fitness.

To determine minimal inhibitory concentration (MIC) before and after evolution, we used the standard qualitative broth microdilution technique (Wiegand et al., 2008). Antibiotic concentrations were serially increased 2-fold until growth inhibition was observed. The lowest concentration at which inhibition was observed was designated the MIC. These results were confirmed by streaking out the glycerol stock on agar plates carrying the relevant antibiotics at MIC as well as the concentration that was 2-fold below the MIC.

### Quantifying fitness costs of altered translation accuracy and antibiotic exposure

To determine the fitness costs arising from antibiotic exposure as well as altered translation accuracy, we measured ancestral (‘anc’) and evolved (‘evo’) growth rates in the presence vs. absence of antibiotic (for antibiotic cost) and relative to ancestral WT in LB (for the cost of altered accuracy, which arises from any deviation from WT translation accuracy) (Figure 1a). Specifically, we used the following calculation for each strain.

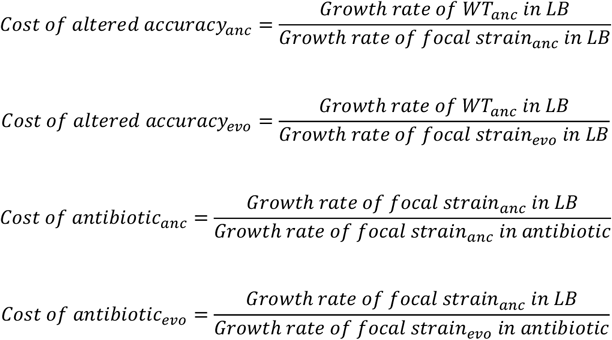

We used ancestral growth rates in LB for the numerator so that as the costs increase, the value of the fraction increases, making it simpler to assess the magnitude of the cost. Note that the antibiotic cost is not a measure of the cost of antibiotic resistance, although in evolved lines both resistance and compensation for resistance are likely to contribute to the antibiotic cost calculation. In principle, this calculation for costs can be applied to any strain, irrespective of its prior exposure to antibiotic. Accuracy costs are defined relative to ancestral WT growth in LB, and will only have biological meaning if applied to strains whose translation accuracy had been altered.

### Whole-genome sequencing to identify mutations and determine mutation spectra

In order to track genotypic changes that occurred during adaptation, we subjected populations from our 15-day stocks (and where required, from other timepoints) to whole genome sequencing. We extracted genomic DNA (GenElute Bacterial Genomic DNA kit, Sigma-Aldrich, and Wizard DNA extraction kit, Promega) and quantified it (Qubit HS dsDNA assay, Invitrogen). We prepared paired-end libraries using the Illumina Nextera XT DNA library preparation kit following the manufacturer’s instructions. We then sequenced all 15-day evolved lines on an Illumina HiSeq 2500 platform using the 2×100 bp paired-end reaction chemistry, and obtained ~3 million reads per sample with quality >Q30 (reads below this quality cut-off were discarded), corresponding to an average per base depth of ~100X. Our WT strain KL16 (a derivative of MG1655 with ~50 mutations) does not have a sequenced genome available. Therefore, for each sample, we aligned quality-filtered reads to the NCBI reference *E. coli* K-12 MG1655 genome (RefSeq accession ID GCA_000005845.2) using the package Breseq (Deatherage and Barrick 2014). After removing ancestral (KL16) mutations from evolved lines, we extracted a list of base-pair substitutions and indels from the Breseq output. We only counted mutations with >50% frequency, and verified that all were represented in at least 10 reads, before proceeding with further analysis. For the MA lines, reads from pooled samples were aligned to the MG1655 genome using the Burrows-Wheeler short-read alignment tool, BWA (Li and Durbin 2009). We generated pileup files using SAMtools (Li et al. 2009) and used the VARSCAN package to extract a list of base-pair substitutions and short indels (< 10bp) (Koboldt et al. 2009). We only retained mutations with >80% frequency that were represented by at least 5 reads for further analysis.

### Evolution under a mutation accumulation (MA) regime to determine mutation rate and spectrum

To test whether altering translation accuracy changes the mutation rate or spectrum, we conducted a mutation accumulation experiment, allowing populations to evolve largely under genetic drift followed by WGS to identify mutations, as described earlier (Sane et al, 2018). We founded 50 MA lines each for the three strains KL16 (WT), KL*ΔZWV* (Mis-1) and KL*ΔZWV*(HA) (Mis-1 + HA). For each line, in order to minimize the impact of selection, we streaked out a randomly selected colony (on or near a pre-marked spot) on a fresh agar plate every 24 h, thereby passing the population through single colony bottlenecks. Every 3 days, we inoculated a part of the transferred colony in LB broth at 37°C for ~3 hours and froze 1 mL of the growing culture with 8% DMSO at −80 °C. For the current study, we only used stocks frozen on the final day, i.e., day 60. Based on previous work (Sane et al., 2018, Sane et al., 2023), we expected this time to be sufficient for the accumulation of one mutation per line on average. We pooled 10 lineages from the last (60 day) timepoint creating 5 pools per strain and extracted DNA from a total of 15 such pools for the 3 strains mentioned above. To estimate mutation rate and spectrum, we considered the total mutations present across all 5 pools (5×10=50 total lines) for a given strain at day 60, and averaged that number across the number of lines. 60 days with 24-hour plate transfers at ~27.5 generations per day corresponds to ~1650 generations. Mutation rate was calculated as:

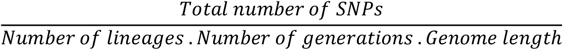

We had 50 lineages, 1500 generations total (~25 in 24 hours) and 4,641,652 bp as genome length. Note that due to the pooling, we could not determine exactly how many mutations occurred in any given line; hence, we could not estimate confidence intervals for mutation rates. To determine the mutation spectrum, we quantified the proportion of different types of observed mutations.

## RESULTS

### All strains with altered translation accuracy show rapid adaptation across regimes

To assess the role of altered translational fidelity in adaptation to antibiotics, we evolved strains with varying levels of mistranslation (~10 to 50 fold different from the WT, (Samhita et al., 2013, Agarwal et al., 2015, Samhita et al., 2020, Stikeleather et al., 2025)) for ~300 generations in one of three antibiotics possessing different modes of action: ciprofloxacin (Cip), kanamycin (Kan) and ampicillin (Amp), and one (control) antibiotic-free regime. We genetically increased wild type (WT) mistranslation rate either by using a strain with depleted initiator tRNA content (‘Mis-1’, same as the ‘Mutant’ in (Samhita et al., 2013)) or by introducing a mutation in the ribosomal protein S4 (‘Mis-2’). To reduce mistranslation, we introduced a mutation in the ribosomal protein S12, which generates ‘hyper-accurate’ ribosomes (strain ‘HA’, same as WT(HA) in (Samhita et al., 2020)). Additionally, we included a strain that carried both a mistranslating mutation as well as the hyper-accurate one (Mis-1+HA (Samhita et al., 2020)).

The mistranslating strains Mis-1 and Mis-1+HA paid an initial fitness cost relative to the WT during growth in rich LB medium, whereas the HA strain did not show a cost of hyper-accuracy **(Fig. 1a)**. Note that despite lower mean fitness compared to WT, large variation across replicates of strain Mis-2 meant that this difference was not significant; thus, Mis-2 did not show a significant ancestral fitness cost of mistranslation **(Fig. 1a)**. On adding antibiotics, all ancestors showed a (further) decline in fitness (**Fig. 1a;** see Table S1 for statistical comparisons). Thus, antibiotic pressure exacerbated the cost of mistranslation, and also imposed an independent cost for non-mistranslating strains (WT and HA). In terms of MIC (minimum inhibitory concentration), the cost of antibiotic was not consistent: in Cip, all strains had the same ancestral MIC values; in Kan, only Mis-1+HA showed a marginally lower MIC (7.5 µg/mL vs 10 µg /mL for all others); and in Amp, both Mis-1 and Mis-1+HA showed lower ancestral MICs (3 and 3.5 µg/mL vs 4 µg/mL for the rest, ~0.75 fold difference) (**Fig. S2**).

Our experimental evolution protocol involved dilution with transfer to fresh media every 12 h (~15 generations) and should therefore select for faster growth rate (**Fig. 1b**). Nearly all strains showed a significant increase in growth rate by 300 generations, excepting WT in Kan and Amp and HA in Amp **(Fig. 1c, Table S1)**. Additionally, all strains showed an increase in MIC values (from sub-MIC to ~10X MIC for Cip, ~25X MIC for Amp and ~5X MIC for Kan, Fig. S2, ancestral vs evolved MIC, paired t-test for all strains, t= 4815 for Cip, t= 84.4 for Kan and t=457.4 for Amp, P<0.001 for all comparisons) with no perceptible across-strain differences **(Fig. S2)**. Note that several strains with altered accuracy reached higher absolute growth rates than the evolved WT strain. Both genotype and antibiotic determined the outcome, independently and via genotype by environment (GxE) interactions (ANOVA: evolved fitness~ strain x antibiotic, F_antibiotic_ =97.86, F_genotype_ = 188.2, F_interaction_ =97.6, P<0.001 in each case). Finally, WT control lines evolved under the antibiotic kasugamycin (Ksg, which reduces translation initiation events without affecting tRNA copy number, serving as a control for Mis-1 in LB (see Methods) also showed higher fitness, with the mean growth rate increasing from 0.53 to 1.37 (WT ancestral vs evolved, t test, t=29.2, P<0.001). Interestingly, Mis-1 evolved in LB reached roughly the same mean value of absolute growth rate, increasing from 1.1 to 1.35 (Mis-1 ancestral vs. evolved, t test, t= 10.84, P<0.001), suggesting that the use of WT+Ksg to mimic Mis-1 growth was appropriate.

At the early stages of the evolution experiments, during the 12 h population transfer we inoculated a second microplate to estimate population level growth rates. Given the lack of replication, we were not confident about the accuracy of these growth rates. Nonetheless, the data suggested that within 4 days of evolution (~80 generations), the mistranslating strains had already increased fitness by ~10 to 25% (data not shown). Population level whole genome sequencing showed that at this early stage, several mistranslating populations harboured mutations in antibiotic resistance genes (e.g., *gyrA, fusA*, and *marR* mutations) at >80% frequency; but these were rarely observed in WT and HA populations **(Table S2)**. These results suggested that mistranslating strains adapted at faster rates, via antibiotic-specific mutations that likely compensated for the fitness costs imposed by each antibiotic.

### The cost of antibiotic (but not accuracy) declines consistently over the course of adaptation

To delineate the costs imposed by mistranslation versus exposure to antibiotic, we assessed fitness costs (**Fig. 1a**) for both ancestral and evolved strains. The cost of antibiotic reflects the proportional reduction in growth rate of the focal strain in the absence vs presence of antibiotic (as a ratio), and the cost of altered accuracy reflects the reduction in growth rate of the WT ancestor vs. the focal strain in LB (see Methods for details). In both cases, the absence of a cost would result in a value of 1, which should occur when there is no perturbation of growth rate by addition of antibiotic or by altering accuracy. Increasing values above 1 would indicate higher costs, whereas values below 1 would indicate a relative benefit. Note that the antibiotic cost here is only a measure of growth rate retardation in the presence of antibiotic, and does not reflect fitness costs associated with mutations that provide antibiotic resistance.

All antibiotic-evolved lines showed a significant decline in the cost of antibiotic exposure (henceforth called cost of antibiotic), except WT and HA in Amp and WT in Kan **(Fig. 2a, Table S1)**. In contrast, the cost of mistranslation changed only across specific genotype-antibiotic combinations **(Fig. 2b, Table S1; also see Fig. 3 for comparison of antibiotic vs. accuracy costs)**. The difference between Mis-1 and Mis-1+HA is striking: although both strains had initially similar costs of mistranslation **(Fig. 1a and Fig. 2b)**, Mis-1 showed comprehensive cost compensation in all antibiotics, whereas Mis-1+HA did so only in Kan where the load of mistranslation would be exacerbated by kanamycin-induced inaccuracies (**Fig. 2b**). Similarly, the difference between strains HA and Mis-2 is also noteworthy. Neither strain had a significant ancestral cost of altered accuracy (**Fig. 1a and 2b**), leading us to expect that accuracy costs would remain unchanged in these strains during adaptation. However, while neither strain showed significant cost compensation in Cip, strain HA showed an increase in accuracy cost when evolved in Amp and Kan, while Mis-2 costs did not change significantly in either antibiotic (**Fig. 2b**). Importantly, all LB-evolved lines reduced the cost of altered accuracy, even in cases (such as HA and Mis-2) when the initial cost was not very high (**Fig. 2c**). Thus, accuracy costs were rapidly compensated in all strains evolved in LB, even leading to a relative benefit of altered accuracy (cost below 1) in Mis-2 and HA (**Fig. 2c**); but under antibiotic selection, this seems to be more difficult. Overall, it appears that the cost of antibiotic imposes a stronger selection pressure than altered accuracy, with the cost of antibiotics ranging from 1.5–5x, and the maximum cost of altered accuracy only reaching ~1.4x (**Fig. 3**).

**Figure 2.**
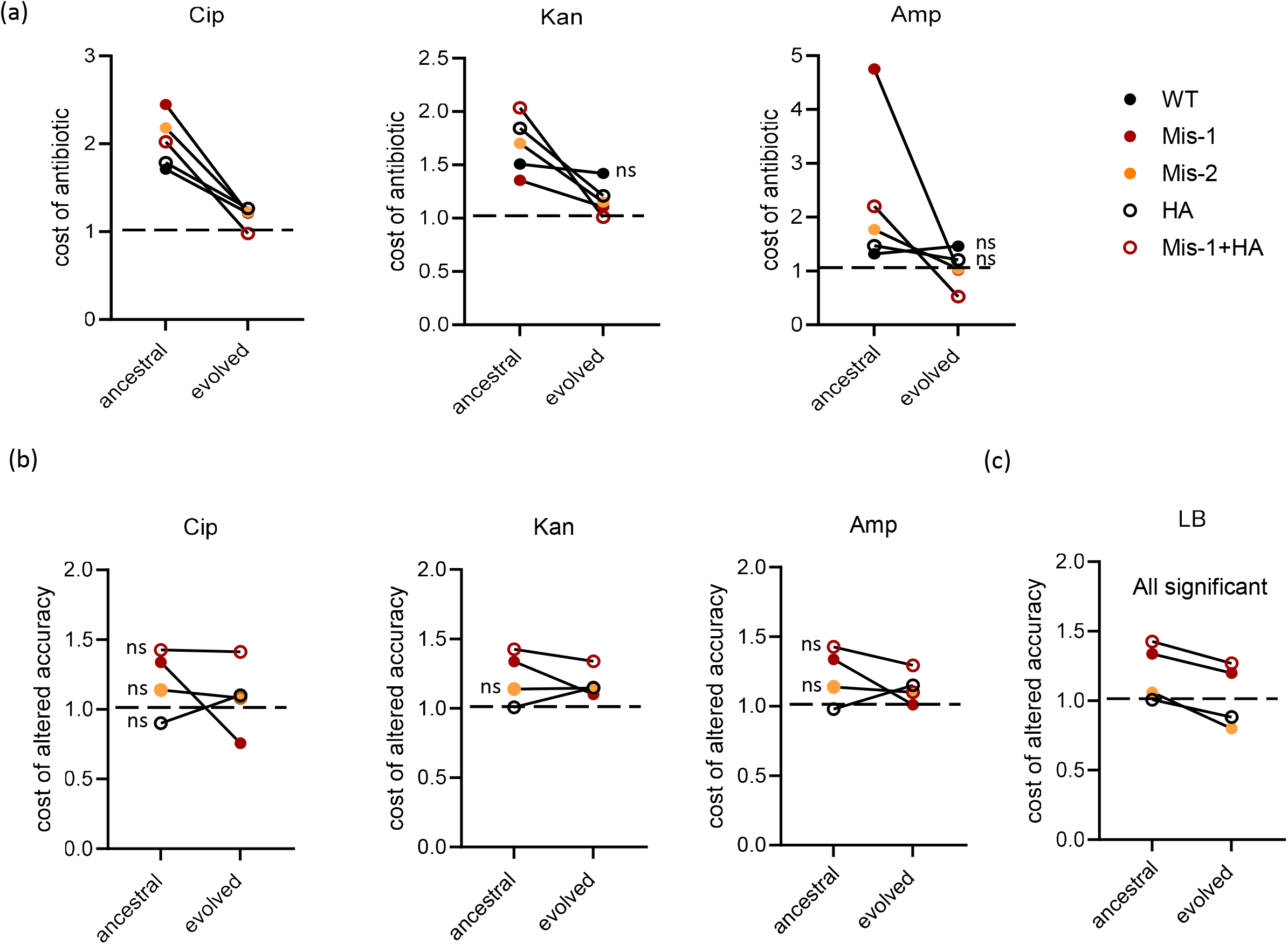
**(a) Cost of antibiotic declines in most strains during adaptation.** Comparison of mean antibiotic costs for ancestral vs evolved lines for a given genotype, tested in the same regimes in which it evolved (n=5 for all except WT and Mis-1 in Kan and Cip, where n=6). All lines showed a significant decrease in cost except when indicated (‘ns’, not significant; Table S1). **(b) Cost of altered accuracy declines for select genotype-antibiotic combinations**. Comparison of mean accuracy costs for ancestral vs evolved lines for a given genotype, tested in the same regimes in which it evolved (n=5 for all except WT and Mis-1 in Kan and Cip, where n=6). All lines showed a significant decrease in cost, except when indicated (‘ns’, not significant; Table S1). Note that HA in Amp showed a significant increase rather than decrease. **(c) Cost of altered accuracy declines for all genotype-antibiotic combinations in LB**. Comparison of mean accuracy costs for ancestral vs evolved lines for a given genotype, tested in the same regime in which it evolved (n=5 for Mis-2 and Mis-1+HA, n=6 for WT and Mis-1 and n=4 for HA). All lines showed a significant decrease in cost, except when indicated (‘ns’, not significant; Table S1).

**Figure 3.**
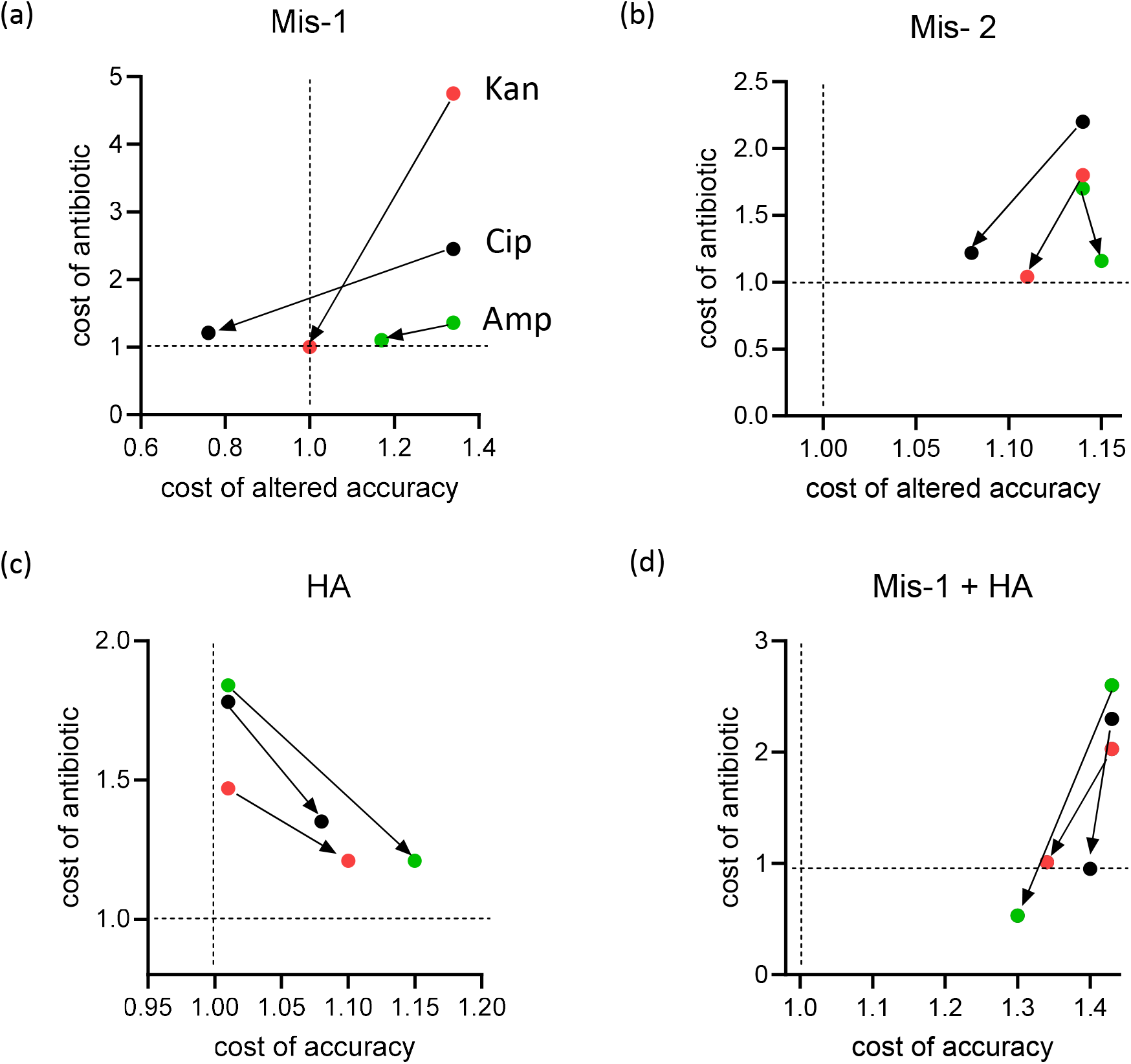
Simultaneous changes in antibiotic and accuracy costs during adaptation. Evolutionary change in cost of antibiotic vs cost of accuracy, for the four genotypes with altered accuracy (n=5 for all except HA in LB, where n=4). Arrows indicate the direction of the change during adaptation. Colours indicate three different antibiotics. Statistical comparisons are detailed in Table S1.

Since we defined the cost of antibiotic by current and not past exposure to antibiotic, we also computed the antibiotic cost for our control lines evolved in LB (**Fig. 4, Table S1**). The antibiotic cost of Cip either increased or did not change (**Fig. 4a**), whereas for Amp and Kan nearly all tested strains showed a reduction in antibiotic cost despite never being exposed to the antibiotic **(Fig. 4b-c)**. Note that in ampicillin, we could only compute growth rates for WT and Mis-1 since all other strains showed noisy growth profiles. Thus, growing in antibiotic-free media for several generations is sufficient for bacteria to compensate for an antibiotic-induced growth defect, despite no direct selection. Similarly, we measured accuracy cost compensation in antibiotic-evolved lines, by testing their growth in LB. Mis-1, which had the highest ancestral accuracy costs, showed cost compensation across all regimes. For the most part, other strains either showed no compensation or, as with HA, increased rather than decreased ancestral accuracy costs after evolution in antibiotics. **(Fig. 4d-f)**. On the other hand, as discussed above, all lines evolved in LB showed a significant compensation of accuracy costs **(Fig. 2c, Table S1)**.

**Figure 4.**
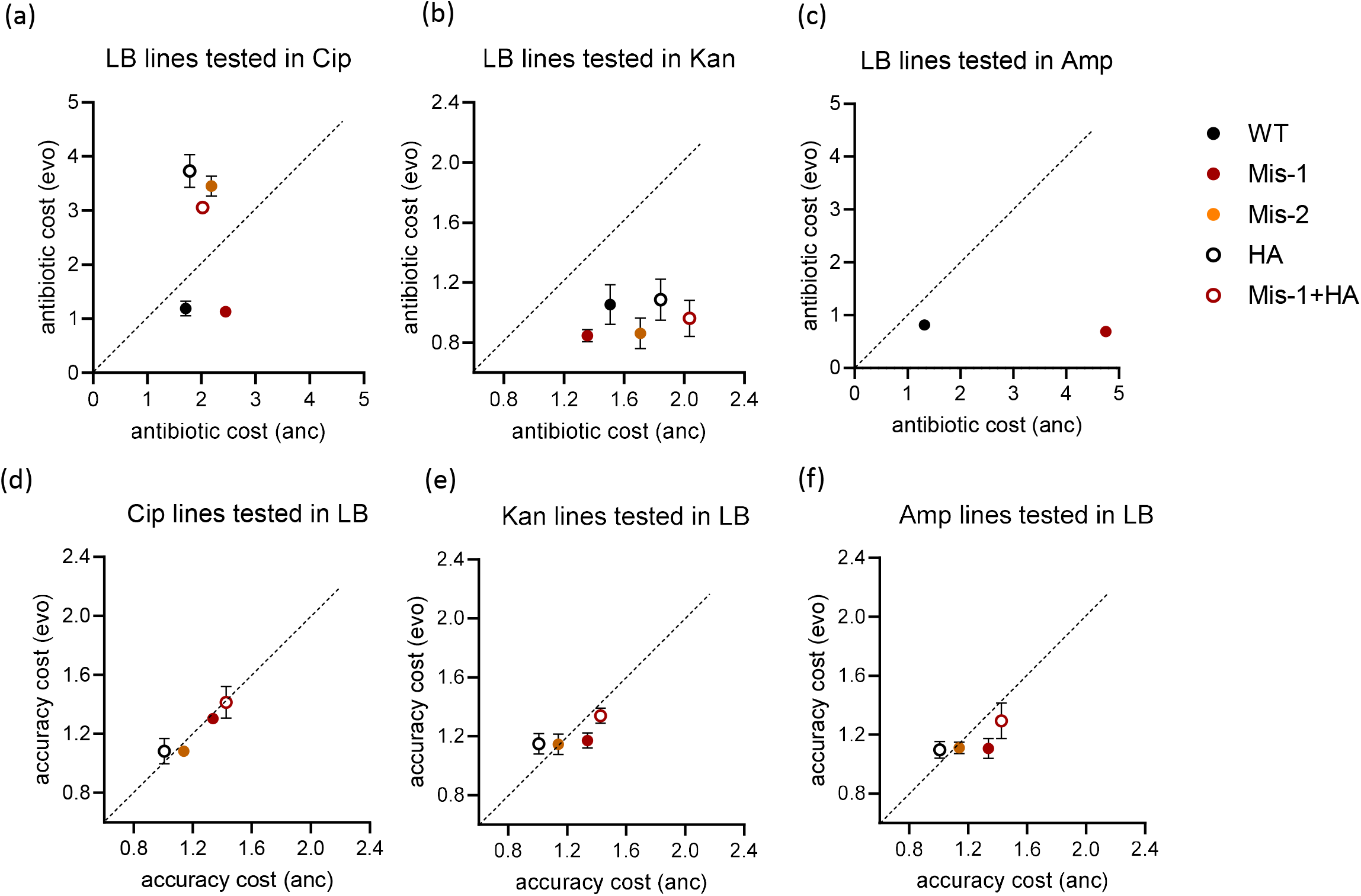
**(a-c) Antibiotic cost compensation occurs in the absence of direct selection.** Mean values (+/−SD) for evolved vs ancestral antibiotic costs across all genotypes for the LB regime (n=5 for all except HA, where n=4). WT and Mis-1 lines evolved in LB were tested at a later time in Cip, and hence ancestral costs for these lines are marginally different from the original ancestral costs, recorded separately in the raw data (Table S6). For Amp, only WT and Mis-1 data could be computed. Statistical comparisons are detailed in Table S1. **(d-f) Cost of altered accuracy remains unchanged in antibiotic-evolved lines**. Mean values (+/−SD) for evolved vs ancestral accuracy costs across all genotypes evolved in antibiotics and tested in LB (n=5 for all except WT and Mis-1 in Kan and Cip where n=6). Values on the diagonal indicate that ancestors and evolved lines have identical costs. Statistical comparisons are detailed in Table S1.

Overall, we found that exposure to antibiotic (at the sub-MIC concentrations used here) extracts a higher fitness cost than altered accuracy (**Fig. 3**) and this cost is compensated in all lines **(Fig. 2a)**, even in strains evolved without antibiotic exposure **(Fig. 4a-c)**. Accuracy cost compensation, on the other hand, occurs consistently only in the absence of antibiotic (**Fig. 2b)**, and is both regime and strain specific.

### Antibiotic target-specific mutations dominate in the adapted populations

To examine the genetic changes that underlie our observations, we subjected all evolved populations from the final timepoint (~300 generations) to whole genome population sequencing. While all lines showed multiple mutations that rose to >50% frequency, genes known to play a role in antibiotic resistance via target alteration or efflux pump activation dominated, particularly in Cip and Kan **(Fig. 5a-c, Tables S3 and S4)**. This agrees with our observation that antibiotic costs impose a strong selection pressure, and are fixed rapidly across lines. However, barring a few shared genes with mutations, all strains with altered accuracy showed distinct mutation profiles in a given antibiotic, indicating that altered accuracy changes the genetic basis of adaptation to a given selection pressure. Notably, Mis-2 populations frequently showed mutations in *eaeH* (with identical nucleotide changes across populations), and *rclA/rclR* (varied intergenic mutations across populations, **Table S3**), including in LB. These mutations may either reflect a strain-specific mutational hotspot, or they may be broadly beneficial mutations that repeatedly rise to high frequency. Prior work suggests that *eaeH* is a cell surface protein (although it could be a pseudogene in K-12 strains (Homma et al., 2002)), and shares sequence similarity with the EaeH virulence factor that aids attachment to intestinal cells in enteropathogenic *E. coli* (Eltai et al., 2020). *rclA* is a flavoprotein that confers protection against oxidative stress (Meredith et al., 2022), while *rclR* is a transcriptional activator in the same regulon (Parker et al., 2013). Thus, it is plausible that these mutations are beneficial under antibiotic selection, specifically in the Mis-2 background. Apart from these mutations, repeatability across biological replicates was generally poor. However, in most cases, non-WT strains acquired high-frequency mutations in a wider range of genes (**Fig. 5a** and **5d**). We find that this applies even if translation accuracy is enhanced rather than reduced.

**Figure 5.**
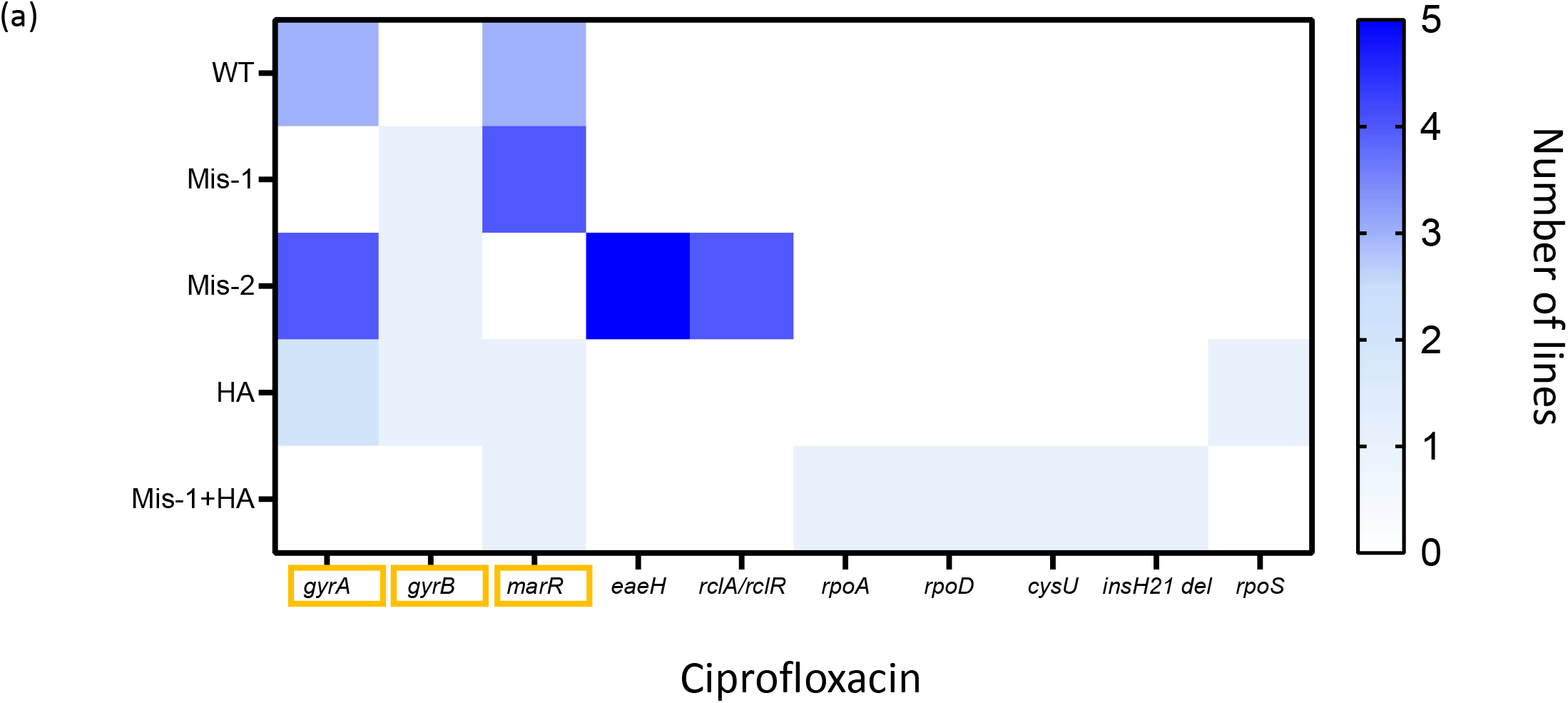
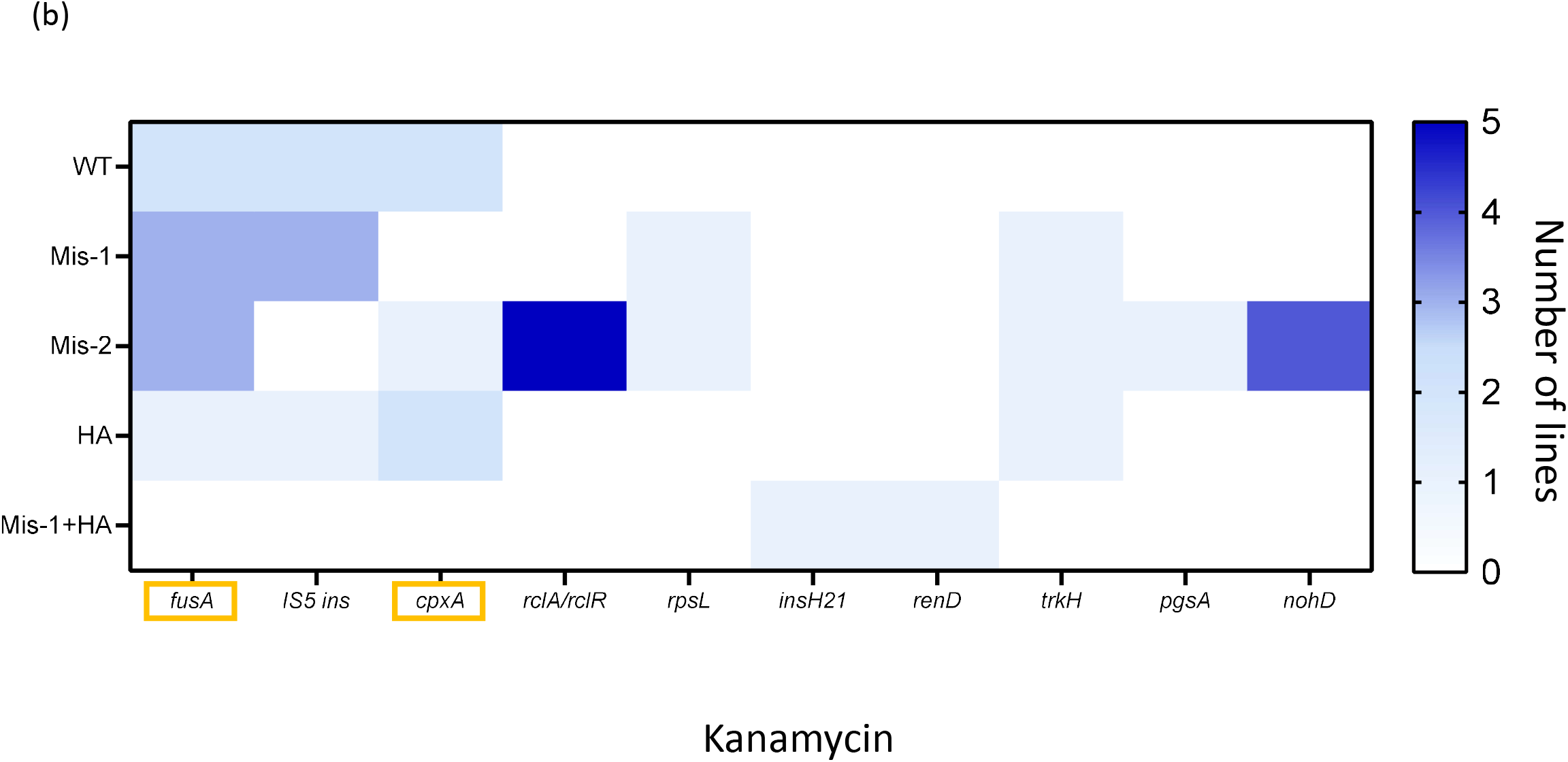
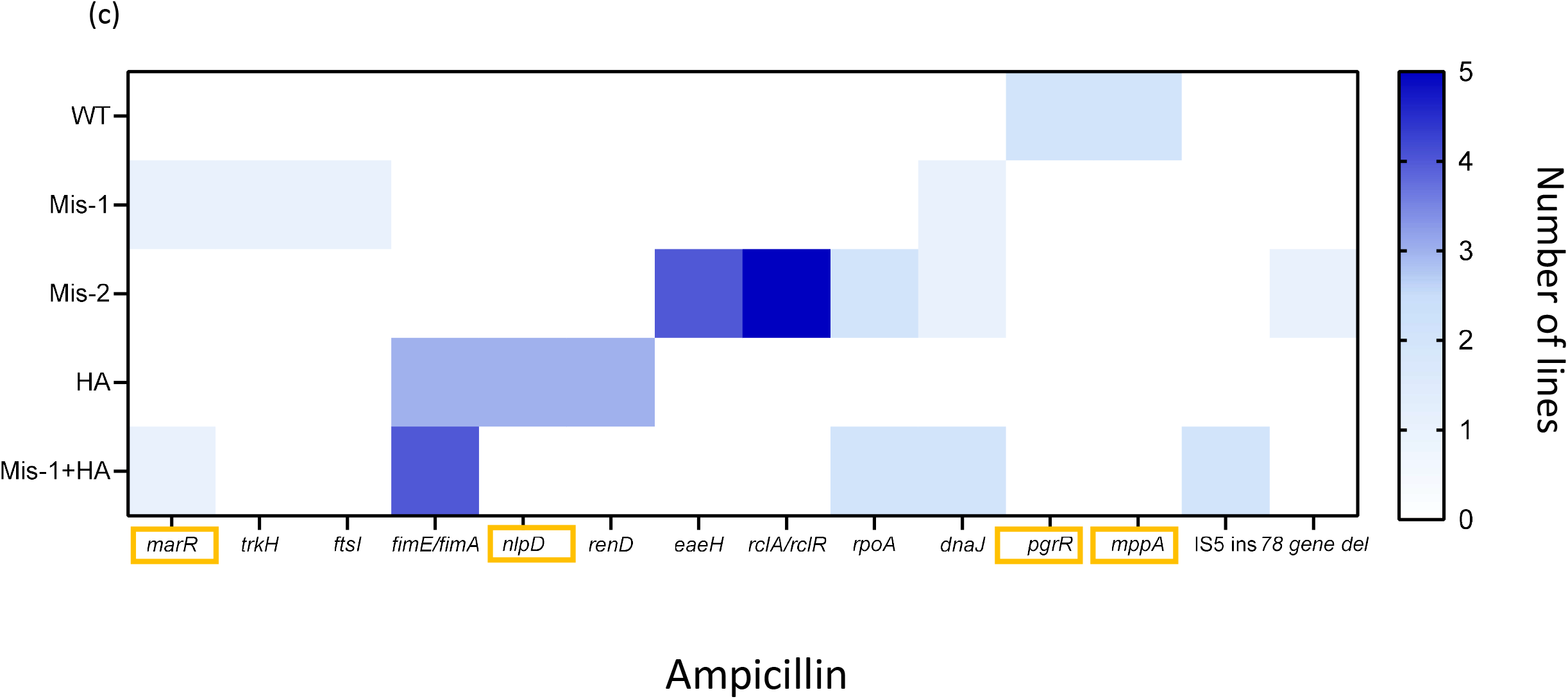
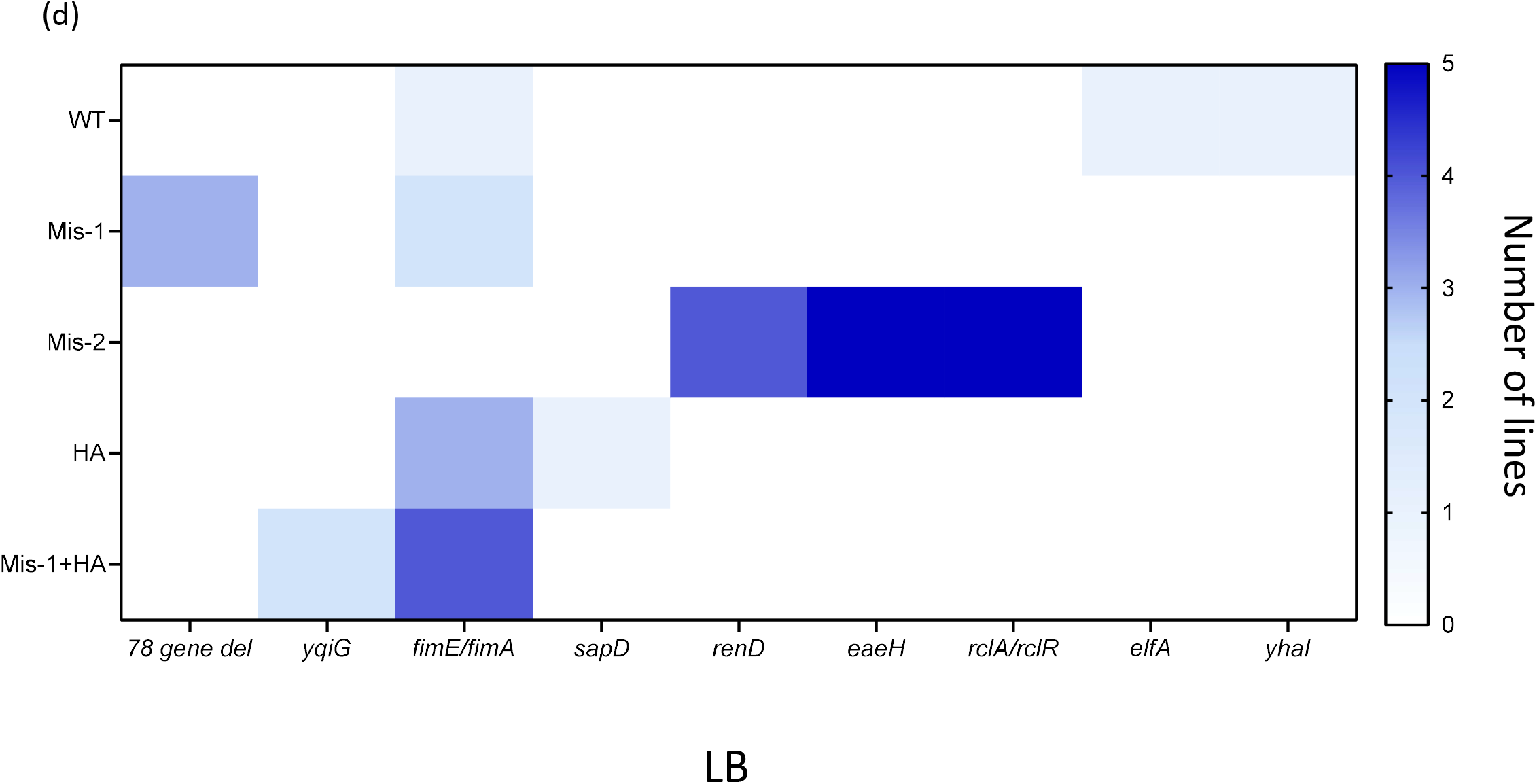
Antibiotic resistance-conferring mutations dominate in evolved lines. Heat maps for all four selection regimes **(a-c)**, antibiotics; **d**, LB, showing gene-level mutations of >50% frequency in evolved populations (n=5 except for HA in LB, where n=4 and WT and Mis-1 in Kan, Cip and LB where n=6). Shades of blue indicate the number of replicate lines that carried a mutation in a given gene; white indicates that no replicates of a given strain showed a mutation in that gene. Known resistance-associated mutations are highlighted in yellow. Details of all mutations are given in Tables S3 and S4.

Next, although the LB-evolved lines all show a significant reduction of accuracy costs **(Fig. 2c)**, none of the high-frequency mutations involve components of protein synthesis **(Fig. 5d)**. It appears therefore that accuracy costs are indirectly compensated, either via epistasis with the original accuracy-altering mutation or through pleiotropic effects of mutations that are not directly connected to protein synthesis. Finally, our experiments included exposing WT to the antibiotic Kasugamycin (Ksg) as a tRNA-independent method of reducing translation initiation events, i.e., a control for our Mis-1 line which lacks three of four initiator tRNA genes. Some evolved mutations in these lines, such as those in RsmA/KsgA, are directly involved in Ksg resistance (Ochi et al., 2009) and all were distinct from Mis-1 in LB (compare **Fig. 5d** with **Fig. S3**). Again, these results emphasize that the mechanism by which protein synthesis is perturbed is important for determining which mutations are fixed, and that antibiotic costs generally dominate over the costs introduced by accuracy changes. Finally, we tested whether mutational diversity is influenced by altered mutation rate or bias in strains with altered translation accuracy. Basal mutation rates and spectra for WT, Mis-1, and Mis-1(HA) estimated by whole-genome sequencing of mutation accumulation (MA) lines were indistinguishable **(Tables 1 and S5**). Thus, the observed mutational diversity is likely to be determined primarily by differential strength and mechanisms of selection imposed by altered accuracy and/or antibiotic exposure.

**Table 1:** Mutation rates and spectra do not change with altered translation accuracy. The observed number of mutations of different types are shown, for each MA-evolved strain. Mutation bias is compared across strains using chi-square tests (e.g., SNP vs. indel bias).

| Mutation type | Strain |  |  | Comparison of mutation bias |  |  |  |
| --- | --- | --- | --- | --- | --- | --- | --- |
|  | WT | Mis-1 | Mis-1+HA | WT vs Mis-1 |  | WT vs (Mis-1+HA) |  |
| | | | | $\chi^2$ | P | $\chi^2$ | P |
| SNP | 64 | 71 | 74 | 0.02 | 0.98 | 0.77 | 0.38 |
| Indel | 8 | 9 | 13 |  |  |  |  |
| Transition | 36 | 39 | 36 | 0.02 | 0.87 | 0.8 | 0.38 |
| Transversion | 28 | 32 | 38 |  |  |  |  |
| AT to GC | 15 | 22 | 23 | 0.74 | 0.39 | 0.95 | 0.33 |
| GC to AT | 34 | 35 | 35 |  |  |  |  |
| Coding | 51 | 50 | 56 | 1.5 | 0.2 | 0.3 | 0.57 |
| Non-coding | 13 | 21 | 18 |  |  |  |  |
| Mutation rate | 1.8 X 10<sup>-10</sup> | 2.5 X 10<sup>-10</sup> | 2.6 X 10<sup>-10</sup> |  |  |  |  |

## DISCUSSION

Mistranslation and its adaptive potential have intrigued biologists for some time now. Recent studies have found that mistranslation can promote adaptation to several inhospitable environments, including during growth in antibiotics. *E. coli* strains show a wide range of translational accuracies (Mikkola and Kurland 1992), so the interplay between antibiotic exposure and mistranslation is likely to be of ecological relevance. However, to properly assess the role of protein synthesis errors in adaptation, the costs imposed by mistranslation as well as the externally imposed selection pressure must be taken into account. In this study, for the first time, we define and quantify the costs of altering translation accuracy and costs incurred by antibiotic exposure, and tease apart their individual impacts on the paths to adaptation. We do so by subjecting strains with both *higher* and *lower* translation accuracy than the WT to experimental evolution for ~300 generations in three bactericidal antibiotics with diverse modes of action, so as to observe the impact of antibiotic-specific effects. We find that antibiotic costs are typically higher than those imposed by altered accuracy, and are always rapidly compensated even in the absence of direct selection; whereas accuracy costs are only compensated occasionally, and most comprehensively in the absence of antibiotics. Our whole genome analysis of cost compensation during evolution shows a good match with the phenotypic data, with mutations that lead to antibiotic resistance dominating in evolved lines. Additionally, altered accuracy changes the genetic basis of adaptation to antibiotics. Although all lines show target mutations, non-WT lines also show some high frequency, strain-specific off-target mutations. Previously, we showed that mistranslation increased phenotypic heterogeneity, and proposed that it could alter evolutionary dynamics across populations (Samhita et al., 2021). In support of this prediction, here we show that altering translation accuracy in either direction can diversify the genetic paths to adaptation under moderate selection pressures (sub-MIC antibiotic concentrations). Given that bacteria in natural environments show a range of mistranslation rates (Mikkola and Kurland 1992), we suggest that simultaneous imposition of antibiotic and translation accuracy costs may shape distinct adaptive outcomes under antibiotic selection.

### Altering translation accuracy in either direction changes the identity of beneficial mutations and increases mutational diversity under moderate selection pressures

Prior work with a reporter gene conferring antibiotic resistance showed that mistranslation increased mutational diversity across replicates (Zheng et al., 2021). Our results both support and extend these data: we find that mutational repeatability varies across strains (i.e., due to altered accuracy), and is also antibiotic-specific. Zheng and colleagues suggested that different replicates of a mistranslating strain may have distinct cellular phenotypes due to idiosyncratic mistranslation events, altering the fitness effect of new mutations and leading to the fixation of different resistance mutations across replicates. But why do we observe this for hyper-accurate strains? It is possible that increased accuracy also imposes hidden costs and generates phenotypic heterogeneity, as observed in mistranslating strains. Although we did not observe a growth cost for our HA strain in this study, it does have a disadvantage when competing against the WT (Samhita et al., 2021), and other hyper-accurate strains are also slower when growing alone (Ruusala et al., 1984). Hyper-accurate ribosomes, while more stringent in decoding, may frameshift more (Sipley and Goldman 1993), and could be losing translation fidelity in other ways. If so, the hypothesis proposed by Zheng et al may well explain our results; but further work is needed to rigorously test the underlying mechanisms.

Among the strains we used, inter-replicate variability was lowest in the WT, where mutations largely occurred in genes encoding known antibiotic targets (e.g.: *gyrA* and *marR* in Cip, *fusA* in Kan and *mppA* and *pgrR* in Amp). When antibiotic regimes were compared, mutations in Cip-evolved lines tended to show higher repeatability, suggesting that paths to Cip resistance may be limited even at sub-MIC antibiotic concentrations. In contrast, mutations in the Amp and Kan regimes were more diverse, both across replicates and across strains. Note that in line with previous work (Pereira et al., 2023), we observed frequent background mutations that drive adaptation to growth in micro-well plates (e.g., mutations in *fimE, fimA, trkH, rpoA, ptsP* and *rpoC*) and likely have no association with antibiotic resistance or translation accuracy. Among the three antibiotics used, we see maximum across-strain mutational diversification in ampicillin **(Fig. 5c)**, and more shared mutations (at the gene level) across strains in Cip and Kan. This suggests that antibiotic modes of action also influence mutational predictability on the background of altered accuracy.

Broadly, strain-specific differences in adaptive mutations must arise from (a) Sampling different sets of mutations (i.e., an underlying mutational bias) (b) Sampling the same set of mutations but fixing different mutations due to direct epistasis with mutations that alter accuracy, or (c) Sampling the same set of mutations but fixing different mutations because of indirect (global) epistasis due to different ancestral fitness as an outcome of altered accuracy. In all cases, pleiotropic effects mediated by the mechanism of mistranslation, mode of action of the antibiotic and the selection pressure (mild vs. moderate vs. high) can also add to mutational differences. Indeed, a different mechanism may be operating in each (or a subset) of our strains, given that the mode of mistranslation influences the degree of phenotypic variability and change in fitness (Samhita et al, Evolution, 2021). More experiments will be needed to distinguish between these possibilities. Changing translation fidelity certainly appears to alter the fitness landscape as predicted by prior models (Schmutzer and Wagner 2023); however, in the absence of a clear relationship between mistranslation level and mutational diversity or evolutionary predictability, we are currently unable to postulate a hypothesis as to how this occurs. It would appear that both the mode of mistranslation and the specific selective pressure (here, the mechanisms of action of specific antibiotics) shape the fitness landscape, altering the mutational basis for adaptation.

### Altered translation accuracy affects multiple aspects of adaptation

Our work adds new dimensions to prior studies implicating mistranslation in altering the rate, degree and genetic basis of adaptation. Early in evolution, high-frequency adaptive mutations are observed only in mistranslating strains, which also appear to adapt faster, commensurate with larger initial antibiotic costs. This observation is similar to earlier work with fungi **(Weil et al**., **2017)** and in line with diminishing adaptive returns (Card et al., 2019). In contrast, the hyper-accurate strain HA with a negligible cost of altered accuracy (i.e., ancestral growth rate comparable to WT) did not show an increase in growth rate or any mutations in this time period. Irrespective of the degree and mode by which translation accuracy was changed, all strains evolved to reach the same MIC values (from sub-MIC to ~10X MIC for Cip, ~25X MIC for Amp and ~5X MIC for Kan, **Fig. S2**). Despite identical final MIC values, strains with altered accuracy often had higher final growth rates than the WT (except in Cip), indicating that altered accuracy may lead to diverse effects on distinct components of fitness. Similar results were reported in earlier work where a plasmid-borne TEM-1 gene for ampicillin resistance was evolved in a WT vs. mistranslating strain background for 6-8 generations under various target antibiotics (Bratulic et al., 2017): the mistranslating strain achieved similar MIC but higher cell densities than the WT in the absence of antibiotic.

### Cost compensation is accompanied by raising but never by lowering translation accuracy

In our work, raising baseline translation accuracy also led to other interesting effects. For instance, while most genotypes adapted by simultaneously reducing antibiotic and accuracy costs, in the strain with hyper-accurate translation the cost of accuracy *increased* during adaptation to antibiotics. Additionally, no mutations relevant for translation accuracy were observed. In contrast, two of the mistranslating lines (Mis-1 replicate 2, and Mis-1+HA replicate 2) that evolved in kanamycin — therefore bearing an increased burden of translation errors — adapted by acquiring mutations that confer hyper-accuracy (H77L and P91Q respectively in the ribosomal protein S12 (Zengel et al., 1977, Siller et al., 2010, Agarwal and O’Connor 2014), likely reducing mistranslation levels. The reason why we sometimes observed mutations that increase accuracy, but none that decrease accuracy, remains unclear and needs further exploration. Overall, it is clear that multiple drivers determine cost compensation for antibiotics and altered accuracy: genotype, magnitude of the cost, mechanism of mistranslation (or hyper-accuracy), and the mode of action of the specific antibiotic.

### Limitations and future directions

In conclusion, our work brings up several new ideas that merit further exploration, while also highlighting limits in our current state of knowledge. An important caveat in our work is that although we define two separate costs, they are necessarily intertwined. We measure antibiotic cost compensation in strains whose translation accuracy is also altered, and vice-versa. We hope that future work will identify other ways of quantifying the costs of accuracy in conjunction with diverse selection pressures. Another caveat is that although we do not see any difference in mutation rates or spectra across strains, higher concentrations of the antibiotics Cip and Amp can enhance mutation rates (reviewed in Andersson and Hughes 2014), and hence, our results may have limited predictive power for adaptive trajectories under high antibiotic concentrations. Finally, while antibiotic costs dominate over costs of altered accuracy, the mechanism of cost compensation for altered accuracy remains a mystery. Measuring translation accuracy in our LB-evolved lines may shed some light on whether the cost compensation occurs through a reversion to WT levels of translation accuracy, or via other unrelated mechanisms.

Despite these limitations, we can now hypothesize that diverse magnitudes of translational accuracy across bacteria (Mikkola and Kurland 1992) may be yet another factor that makes it so difficult to predict the fitness costs of antibiotic resistance (Hinz et al., 2024). Importantly, we find that growth in rich media *alone* for several generations can result in low-level antibiotic resistance with no known target mutations, an observation that does not bode well for the continuing rise of drug resistance. Our results emphasize the role of non-target mutations (reviewed in Hughes and Andersson 2017) that may confer antibiotic resistance through epistasis with the genetic background, via pleiotropic effects, or through constant exposure to sub-MIC antibiotics, activating stress responses (reviewed in Andersson and Hughes 2014). These observations also have implications for clinical treatment, suggesting that if left unchecked, even drug-sensitive pathogens could respond poorly to late antibiotic treatment. We see mutational diversity even across WT replicate lines, suggesting that in addition to potentiating large effect mutations (Pereira et al., 2023), sub-MIC antibiotic exposure may reduce evolutionary repeatability, in our case exacerbated by changes in translation accuracy. This hypothesis needs further exploration, and is especially important given that several antibiotics target protein synthesis. More generally, while the phenotypic effects of mistranslation continue to be explored, we show here that altering wild type translation accuracy in either direction can have significant consequences on bacterial adaptation to external selective pressures. Thus, to better understand and predict bacterial evolution, we suggest that it is important to disentangle, define, and quantify the (internal) costs of altering translation accuracy and the (external) costs imposed by the environment.

## ACKNOWLEDGEMENTS

We thank DA and LS lab members and Vidyanand Nanjundiah for critical comments on the manuscript, and Shazia Parveen for help with running growth curves. We acknowledge funding and support from the Wellcome Trust/DBT India Alliance (grant IA/E/14/1/501771 to LS); Axis bank (support for AB and grant to LS), IISER Pune (fellowship to ST), the Science and Engineering Research Board (SERB, grant SRG/2023/001817 to LS), the National Centre for Biological Sciences (NCBS–TIFR) and the Department of Atomic Energy, Government of India (Project Identification No. RTI 4006 to DA), and the National Centre for Biological Sciences and the Trivedi School of Biosciences (TSB) at Ashoka university for infrastructure and research support.

## AUTHOR CONTRIBUTIONS

LS and DA conceived of the project and designed work; LS, ST, JM and AB conducted experiments; LS carried out analyses with help from JM and AB; LS and DA acquired funding and wrote the manuscript.

